# NMDA-receptor antibodies alter cortical microcircuit dynamics

**DOI:** 10.1101/160309

**Authors:** RE Rosch, S Wright, G Cooray, M Papadopoulou, S Goyal, M Lim, A Vincent, AL Upton, T Baldeweg, KJ Friston

**Affiliations:** Wellcome Trust Centre for Neuroimaging, Institute of Neurology, University College London, UK; Developmental Neurosciences Programme, Great Ormond Street Institute of Child Health, University College London, UK; Department of Pharmacology, Aston University, Birmingham, UK; Department of Paediatric Neurology, Birmingham Children’s Hospital, UK; Department of Clinical Neurophysiology, Karolinska Institute, Stockholm, Sweden; ToNIC Toulouse Neuroimaging Center, INSERM-UT3, University of Toulouse, France; Department of Clinical Neurophysiology, Evelina London Children’s Hospital, Guy’s and St. Thomas’ NHS Foundation Trust, London, UK; Department of Paediatric Neurology, Evelina London Children’s Hospital, Guy’s and St. Thomas’ NHS Foundation Trust, London, UK; Nuffield Department of Clinical Neurosciences, John Radcliffe Hospital, University of Oxford, UK; Department of Physiology, Anatomy and Genetics, University of Oxford, UK.

**Author notes:** Corresponding Author: Richard E Rosch, BMBCh Wellcome Trust Centre for Neuroimaging12 Queen Square London WC1N 3BG United Kingdom. **Funding:** This study was funded by the Wellcome Trust (RER: 106556/Z/14/Z, KJF: 088130/Z/09/Z). **Author contributions**: RER, GC, MP, SW conceived and planned the study. SW, LU, AV, KH designed and conducted the mouse experiments. ML, AV analysed clinical cases. RER, GC, TB, KJF planned and conducted computational analysis of the data. RER, GC, SW wrote the first draft of the manuscript. LU, AV, KH, ML, AV, TB, KJF critically reviewed and edited the final manuscript. **Competing Interests**: A.V. and the University of Oxford hold patents and receive royalties and payments for antibody assays. **Data and materials availability:** Data and code used in this analysis is provided online for open access under https://github.com/roschkoenig/NMDAR_Encephalitis. **Author contributions** SW KH AB LAU designed and conducted the mouse experiments. ML SG GC RER collected, reviewed and classified clinical EEG recordings. GC MP KJF TB RER planned the analysis and modelling of the data. GC RER conducted the data analysis and computational modelling. RER SW GC TB wrote the first draft of the manuscript. RER SW GC MP SG ML KH AV LAU TB TJF edited and reviewed the final draft of the manuscript.

## Abstract

NMDA-receptor antibodies (NMDAR-Ab) cause an autoimmune encephalitis with a diverse range of electroencephalographic (EEG) abnormalities. NMDAR-Ab are believed to disrupt receptor function, but how blocking this excitatory neurotransmitter can lead to paroxysmal EEG abnormalities – or even seizures – is poorly understood. Here, we show that NMDAR-Ab change intrinsic cortical connections and neuronal population dynamics to alter the spectral composition of spontaneous EEG activity, and predispose to paroxysmal EEG abnormalities. Based on local field potential recordings in a mouse model, we first validate a dynamic causal model of NMDAR-Ab effects on cortical microcircuitry. Using this model, we then identify the key synaptic parameters that best explain EEG paroxysms in paediatric patients with NMDAR-Ab encephalitis. Finally, we use the mouse model to show that NMDAR-Ab-related changes render microcircuitry critically susceptible to overt EEG paroxysms, when these key parameters are changed. These findings offer mechanistic insights into circuit-level dysfunction induced by NMDAR-Ab.

## Introduction

Molecular disruptions of synaptic function have recently emerged as important causes of neurological disorders ^1^. For example, antibodies to N-methyl-D-aspartate receptors (NMDAR-Ab) have been identified as an important, treatable cause of autoimmune encephalitis ^2^, with a particularly high incidence in children who make up approximately 40% of patients. Patients with NMDAR-Ab encephalitis show a diverse range of associated symptoms; including behavioural changes, movement disorders and seizures ^3, 4^. Important aspects of the clinical presentation are electroencephalography (EEG) abnormalities, which have been reported in up to 90% of patients undergoing EEG monitoring; between 20-60% of patients also have overt epileptiform discharges or electrographic seizures ^5, 6^. While there are EEG features that are thought to be relatively specific for NMDAR-Ab encephalitis (e.g. extreme delta brush) ^6^, the more common EEG abnormalities are diverse and non-specific, with global abnormalities broadly associated with more severe disease ^7, 8^.

At the whole organism level, NMDAR-Ab cause an increased seizure susceptibility: Passive transfer of patient immunoglobulin (IgG) containing NMDAR-Ab into a mouse model causes increased susceptibility to chemically induced seizures ^9^. NMDAR-Ab directly affect glutamate transmission through reversible loss of NMDARs, resulting in a reduction of miniature excitatory post synaptic currents (mEPSCs) in brain slices ^10, 11^. NMDAR hypofunction is also a hallmark of psychiatric conditions, such as schizophrenia and acute psychosis ^12, 13^ – whose clinical features resemble the neuropsychiatric and behavioural symptoms also seen in NMDAR-Ab encephalitis.

Linking NMDA receptor hypofunction at the cellular level, and a predisposition to seizures at the systemic scale is challenging. In the simplified view of epileptic seizures as a consequence of excitation-inhibition imbalance ^14^, one would expect NMDAR hypofunction to be associated with a reduction of excitation, thus a decrease in seizure susceptibility. Yet at the level of neuronal ensembles, synaptic molecular changes may have a multitude of different emergent effects depending on their effects on both excitatory and inhibitory components of the neuronal circuit. In relation to NMDAR, observations in a range of experimental models motivate several mechanistic hypotheses explaining the effects of NMDAR hypofunction: These include (i) altered excitatory dynamics with a reduction in late excitatory post-synaptic potential components ^11^; (ii) secondary neurotoxicity, reducing the number of functional excitatory connections ^15^; (iii) a reduction of cortical inhibitory interneuron activity ^16^. Furthermore, paradoxical changes in excitatory and inhibitory transmission – resulting from maladaptive homeostatic changes – have been proposed as underlying NMDAR-Ab associated abnormalities at different temporal scales ^10^.

Relating observations of pathological brain dynamics to these specific hypotheses is challenging. In a highly non-linear dynamic system, such as the brain, the link between synaptic abnormalities and whole brain responses is seldom intuitive or predictable. Neuronal systems are hierarchically structured and each observational scale constrained by larger scale processes, as well as interacting with emergent properties arising from smaller scales ^17^. Some of these multi-scale dynamics can be successfully captured in computational models of neural population dynamics, which have been integrated into validated analytic frameworks, such as dynamic causal modelling (DCM) ^18–20^.

DCM rests on ‘mesoscale’ neural mass models that capture the average behaviours of neural populations at roughly the scale of a cortical column – the model used here is representative of generic features of layered cortical architectures and is often referred to as the canonical microcircuit or CMC ^21^. The parameters of these models (which describe features such as synaptic connection strengths, and population response dynamics) can be fitted to macroscale neurophysiological recordings such as EEG, or LFP recordings and competing models can be ranked according to their model evidence.

Here, we report the results of a dynamic causal modelling analysis of (a) changes in spontaneous (resting state) activity in a subacute mouse model of NMDAR-Ab encephalitis, and (b) abnormal EEG paroxysms observed in a series of paediatric patients with established NMDAR-Ab encephalitis. In a two-stage analysis, we first model the NMDAR-Ab effect in the mouse model; using DCM to identify a minimal set of synaptic parameters required to produce the NMDAR-Ab effect on observed LFP recordings. Using estimates of fluctuations in the parameters identified – based on patient EEG data – we then asked whether the presence of NMDAR-Ab alters network responses to estimated parameter fluctuations. This modelling integrates evidence from patients and experimental animals and provides direct evidence linking microscale observations on NMDAR-Ab-associated dysfunction with dynamic brain phenotypes. This approach enabled us to ask whether NMDAR-Ab-associated cortical dysfunction can be explained by changes in inhibitory or excitatory cortical coupling, or the synaptic dynamics of cortical transmission. Furthermore, we were able to test *in silico* whether (a) paroxysmal abnormalities seen in human patients can be explained by intrinsic, normal fluctuations in cortical coupling – on the background of pathological cortical microcircuitry – or (b) whether they are representative of pathological fluctuations *per se.* This distinction may have implications in terms of the appropriate treatment of paroxysmal EEG abnormalities – either through resetting the system, or controlling abnormal fluctuations.

## Results

### NMDAR-Ab alter the dynamic response to acute chemoconvulsants in mice

Cortical dysfunction associated with NMDAR-Ab was tested in C57BL/6 mice using a two-by-two design. This design tested for the effects of NMDAR-Ab (delivered via intracerebroventricular injection), the acute chemoconvulsant pentylenetetrazole (PTZ, delivered via a later intraperitoneal injection), and their interaction. LFPs were recorded wirelessly in freely behaving animals and 45 minutes of recordings pre- and post PTZ injection of 8 NMDAR-Ab positive and 5 control animals were included for the analysis reported here.

Antibodies alone caused a moderate suppression of the LFP signal across low frequency bands (delta and theta range) in the NMDAR-Ab positive mice. However, additional exposure to PTZ revealed a marked difference between NMDAR-Ab positive and control mice, with a large increase of low frequency (delta-band, 1-4Hz) power in the antibody positive treated mice only (Fig 1). Analysis of variance (ANOVA) revealed a significant main effect of NMDAR-Ab on log-delta-band power [F(1,4601) = 9.67; p = 0.002]; and a significant interaction between NMDAR-Ab and PTZ exposure [F(1,4061) = 85.05; p < 0.001]. A PTZ-induced increase in paroxysmal fast activity consistent with epileptic seizures was observed in the NMDAR-Ab positive IgG treated mice compared to control animals, which has previously been reported elsewhere ^9^. An example of the induced, non-epileptiform slow activity is seen in the bottom panel of Fig 1B. These slow wave cortical dynamic abnormalities were further analysed in the modelling below.

**Figure 1.**
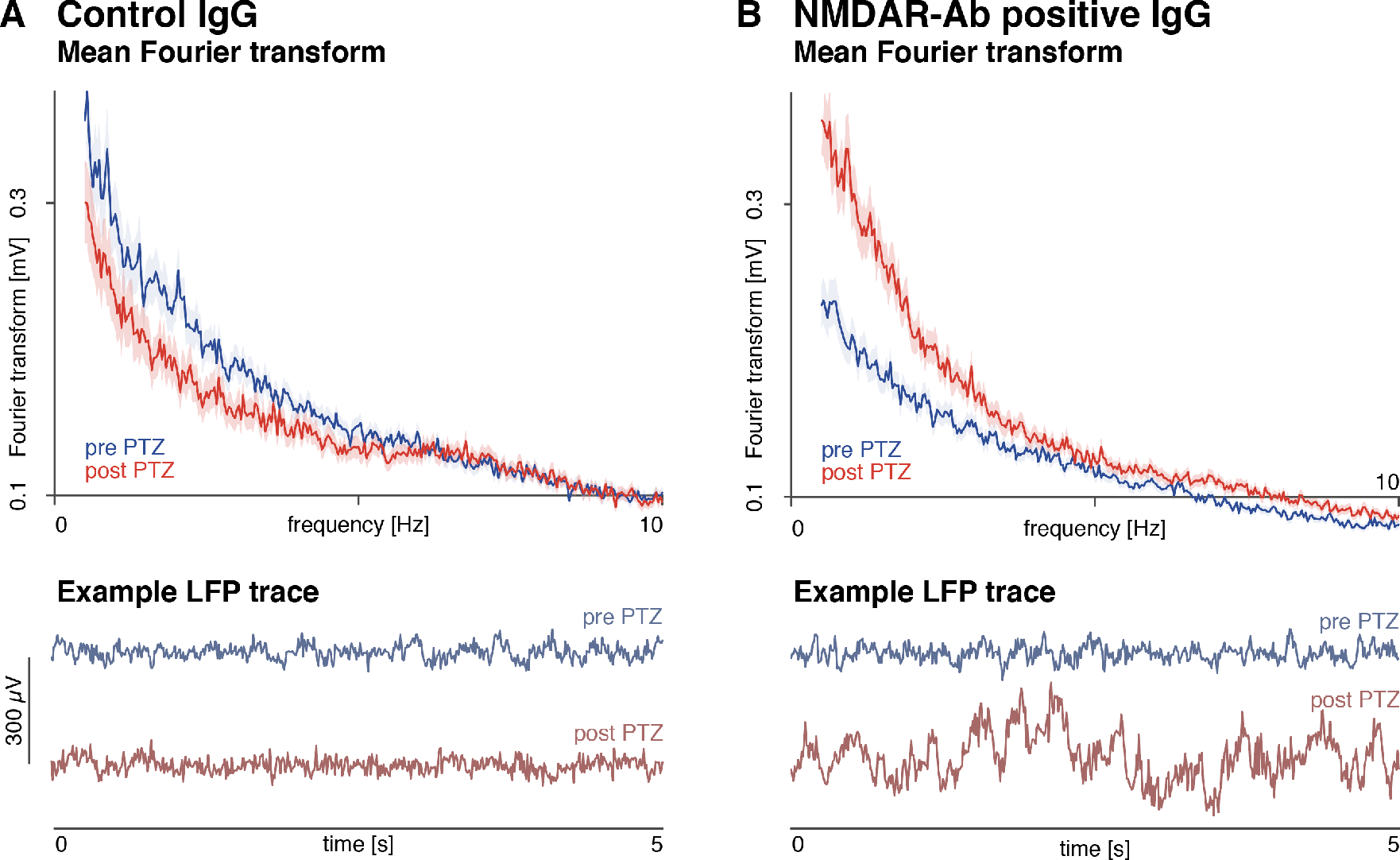
NMDAR-Ab alter the spectral composition of resting state activity following PTZ. Average Fourier spectra of LFP recordings of endogenous activity in mice are shown. (A) In control animals, PTZ injections cause a small decrease in low frequency power. (B) In NMDAR-Ab positive IgG treated animals, PTZ causes a profound increase in low frequency power, which is also visible as high power slow waves in segments; largely without overt epileptiform activity (example shown). Average Fourier spectra across animals are shown for 45 minute recordings pre- and post-PTZ injections, shading indicates the 95% confidence interval. Example 5s LFP segments are also shown for individual animals pre- and post-PTZ injections.

### NMDAR-Ab potentiate PTZ-induced effects in cortical microcircuitry in mice

To explain the observed differences in spontaneous activity, a hierarchical dynamic causal model was used to infer parameter changes associated with the experimental variables over time (i.e. NMDAR-Ab exposure, PTZ infusion, Antibody-PTZ interaction). In brief, a sliding window (length = 30s, step size = 15s) was used to estimate the mean power-spectra over successive time points. Each time window was then modelled as the steady state output of a canonical microcircuit (CMC) model ^21^ with fixed synaptic parameters. By repeating this analysis over windows, we could then identify fluctuations in synaptic parameters that corresponded to the experimental interventions. Across windows, the evolution of spectral patterns was captured well for all experimental conditions (Fig 2A-B). To infer experimental effects associated changes in the DCM parameters, the sequence of parameter estimates was then modelled using a parametric empirical Bayesian (PEB) approach ^22^. Here, slow fluctuations of cortical coupling were modelled as between-window changes in the synaptic parameters estimated within-window (see Papadopoulou et al. ^23^ for a worked example). We included three main experimental effects of interest: (a) NMDAR-Ab, (b) PTZ, and (c) an NMDAR-Ab x PTZ interaction term (Fig 2C).

**Figure 2.**
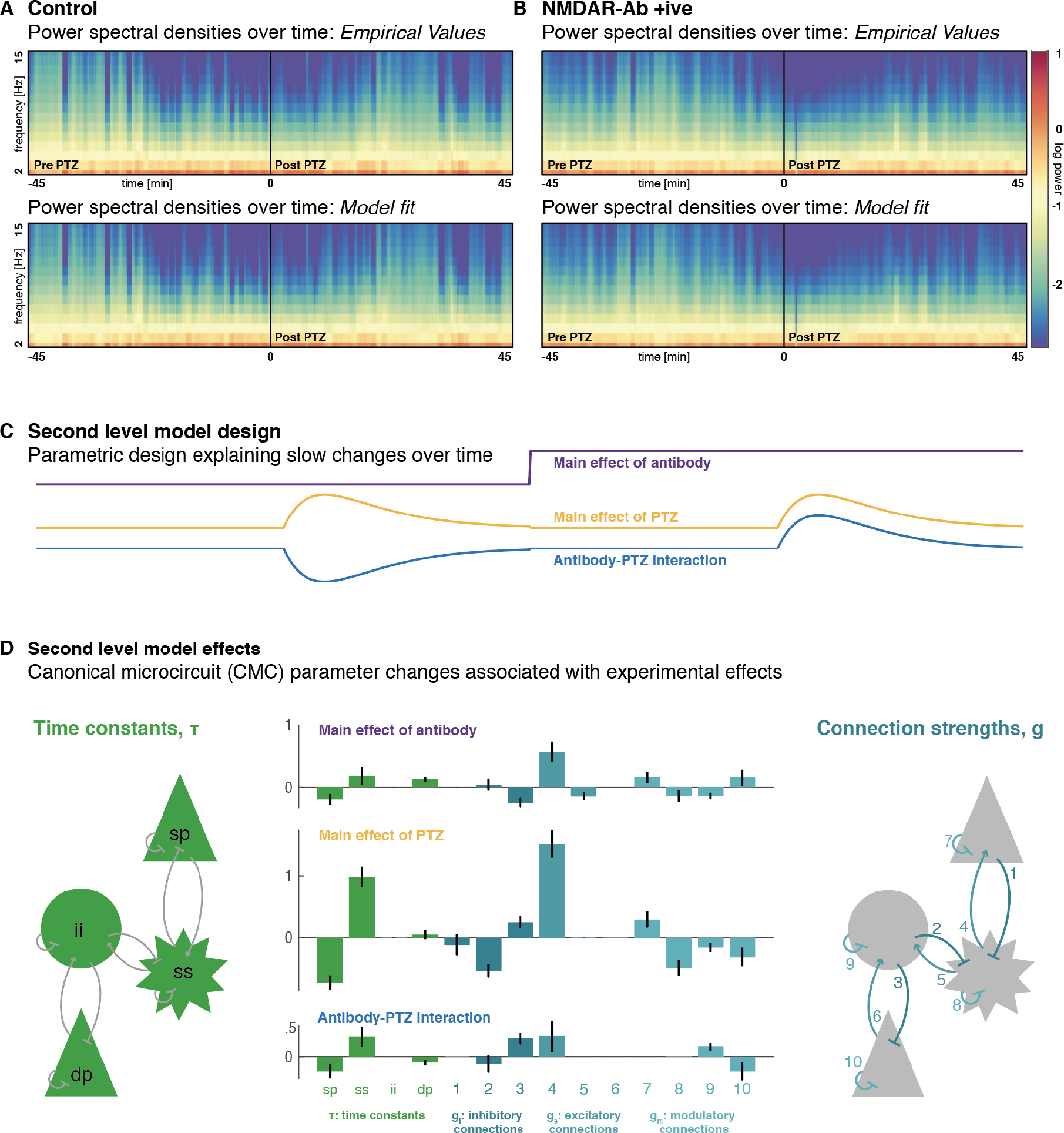
Synergistic changes in synaptic coupling explain the effects of PTZ and NMDAR-Ab. (A,B) DCMs were fitted to sliding window power spectral density summaries of LFP recordings separately for control, and NMDAR-Ab positive animals. Top panels show the observed power spectra over time, with model fits shown in the lower panels. (C) A second-level general linear model with was used to estimate parameter changes associated with NMDAR-Ab exposure, PTZ, and their interaction. The regressors for the three main effects are shown. (D) These experimental effects are associated with parameter changes across all populations of the canonical microcircuit (CMC) neural mass model. The left panel illustrates the population specific synaptic time constants that parameterise the temporal dynamics of post-synaptic responses within that population. The right panel indicates the connections between populations, which are excitatory, inhibitory connections between populations, or self-inhibitory connections. The centre panel shows how each of the parameters is modulated by each of the experimental effects. The strongest effects are caused by PTZ, with the biggest associated changes in *sp* and ss time constants and excitatory connection strength 4. These changes are further potentiated by NMDAR-Ab exposure. Error bars indicate Bayesian 95% confidence intervals. sp: superficial pyramidal cells, ss: spiny stellate cells, *ii*: inhibitory interneurons, *dp*: deep pyramidal cells

The neuronal parameters that affect the spectral composition of spontaneous neuronal activity correspond roughly to the mechanistic hypotheses outlined above: (i) time constants of the neuronal populations *τ* describe the dynamics of neuronal population responses; (ii) excitatory coupling parameters *g_e_* describe the strength of excitatory between-population connections; (iii) inhibitory coupling parameters *g_i_* represent the strength of inhibitory between-population connections, whilst modulatory coupling parameters *g_m_* represent the strength of inhibitory self-connections ^21^.

Spectral changes associated with NMDAR-Ab, PTZ exposure and the interaction were each associated with several parameter changes. The biggest effects were associated with PTZ exposure, with a decrease in the superficial pyramidal cell population time constant (i.e. a faster return to baseline after perturbation), an increase in the spiny stellate population time constant (i.e. a slower return to baseline after perturbation), and an increase in the excitatory connectivity from spiny stellate to superficial pyramidal cells. Notably those changes were further potentiated by NMDAR-Ab and the NMDAR-Ab x PTZ interaction (Fig 2D).

### Shifts in synaptic dynamics underlie emergence of low frequency power in mice

We further investigated the effect of changes in synaptic parameters on the main spectral data feature of interest (i.e. delta-band power). For this, we first performed a principal component analysis over the slow (between time-window) fluctuations of time constants (Fig 3A) and connection strengths (Fig 3B), retaining the first principal component of each. This analysis showed that most of the variance over time can be explained by fluctuations in a small subset of parameters; specifically, the time constants of superficial pyramidal and spiny stellate cells, and the excitatory coupling between them (as is apparent in the analysis in Fig 2).

**Fig 3.**
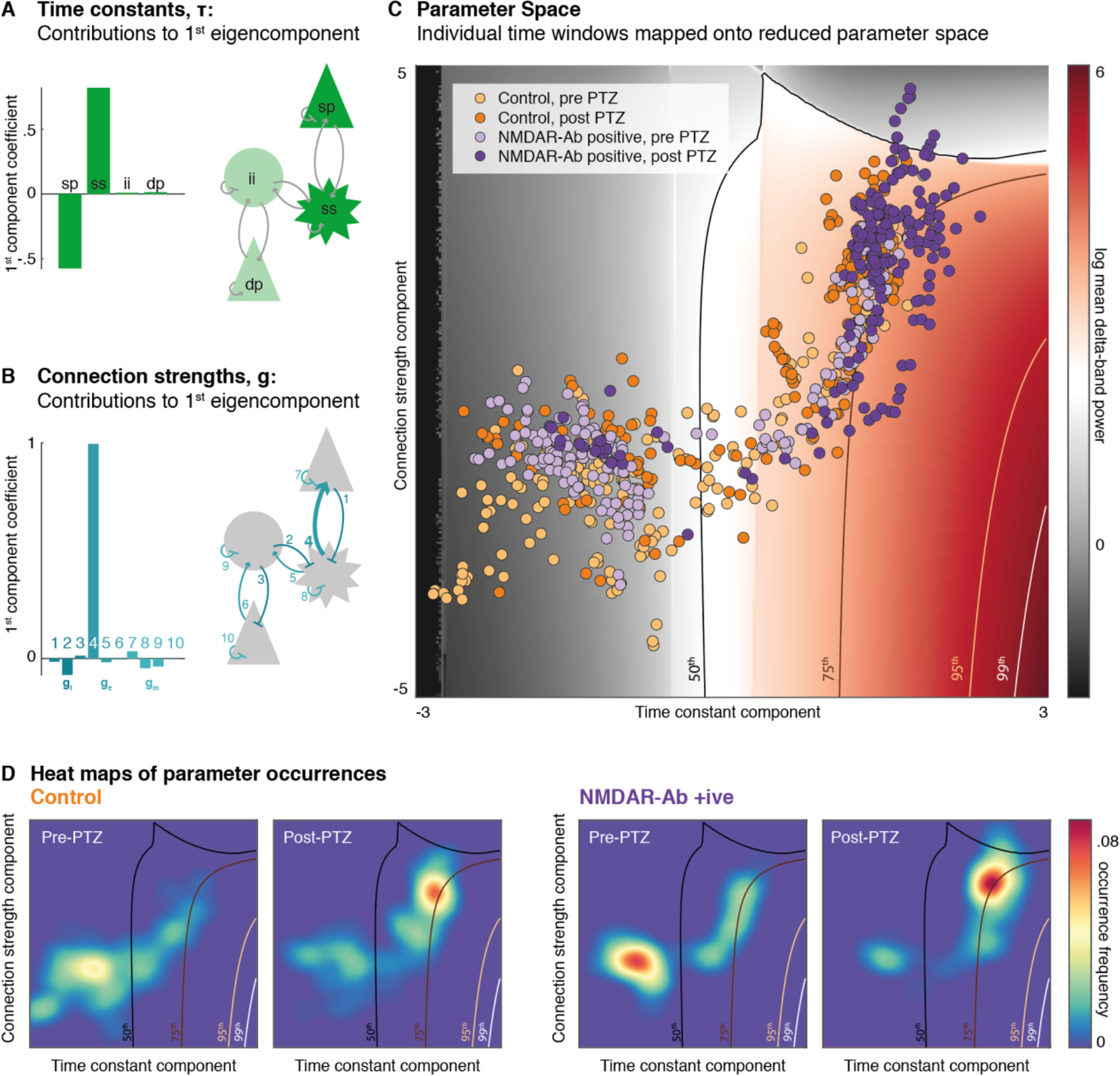
NMDAR-Ab push the neuronal ensemble into high delta-band power regions of reduced parameter space. Parameter variations between time windows are projected onto the first principal component of (A) time constant changes consisting predominantly of superficial pyramidal cell, and spiny stellate cell changes, and (B) of connectivity strength changes consisting predominantly of spiny stellate to superficial pyramidal cell coupling changes. (C) Across this parameter space, simulations can predict spectral densities, of which log mean delta power is shown here (with selected centile isoclines shown). Individual time windows across the four conditions are then projected into the same reduced parameter space, showing an accumulation of NMDAR-Ab positive, post PTZ time window estimates in high delta ranges. (D) The distribution of time windows in parameter space is further illustrated with smoothed heat maps of parameter combination occurrence frequencies over the same section of parameter space for control animals (left) and NMDAR-Ab positive animals (right). Estimates in NMDAR-Ab positive animals cross the 75^th^ centile more frequently than in controls.

We use these two components to project synaptic parameter estimates at each point in time (i.e. window) onto the two dimensions explaining most of the variance (i.e. one time constant component and one connection strength component). To characterise different locations in this parameter space – in terms of the neuronal dynamics generated by the parameters – we used the mean delta band power. This functional characterisation of parameter space is shown (in log-scale) with a colour code and as isoclines indicating mean delta-band power centiles: see Fig 3C. Whilst there is variation in delta-band power associated with both the time constant (x-axis) and the connection strength (y-axis) parameters, the time constants have the greatest effect on delta power: The difference between controls and NMDAR-Ab positive animals in the delta-band power post-PTZ is largely conferred by shifting the time constant component, causing it to cross the 75^th^ delta-band power centile much more frequently than in controls (Fig 3D). This differential effect of PTZ can be seen clearly by comparing the orange and purple dots in Fig 3C.

### EEG paroxysms in patients are caused by fluctuations in synaptic dynamics

To identify which synaptic parameters cause paroxysmal EEG abnormalities commonly observed in NMDAR-Ab encephalitis, we used the above CMC model to perform a DCM analysis of 8 paediatric cases; for which EEG recordings were available and contained visually apparent EEG paroxysms. Briefly, routine visual EEG analysis was performed to identify paroxysmal abnormalities by two EEG-trained clinicians (RER, GC, see Tab 1). For each patient, 2s time windows containing spontaneous activity, short isolated paroxysms, or rhythmic / ongoing epileptiform activity were extracted and used for further analysis (Fig 4A)

**Figure 4.**
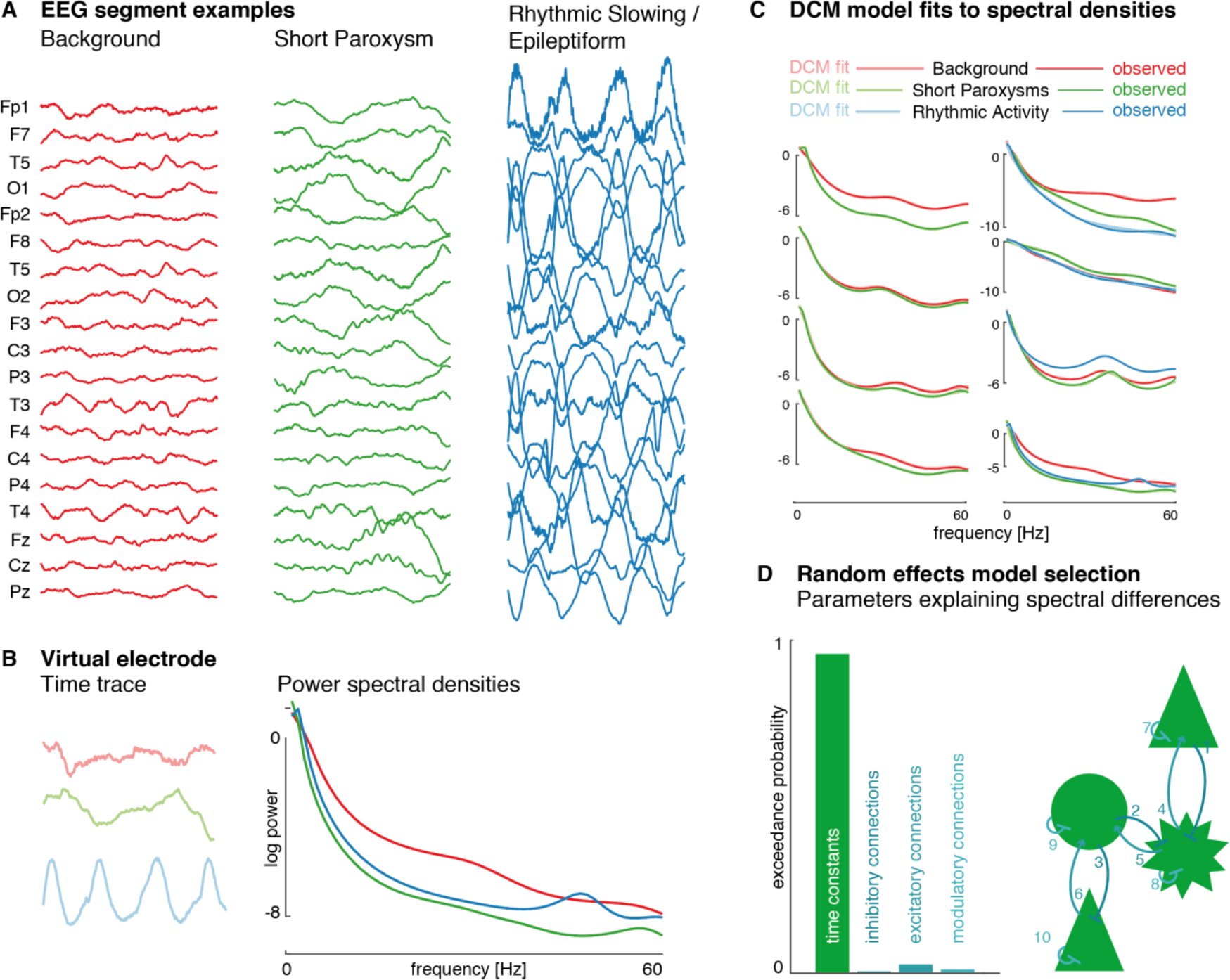
EEG paroxysms in NMDAR-Ab encephalitis patients are best explained as time constant fluctuations. (A) For each individual patient, 2s time windows containing spontaneous activity, short EEG paroxysms and where available longer rhythmic EEG activity were extracted. (B) These were source localised and ‘virtual electrode' time traces extracted at the estimated cortical source. Normalised power spectral density averages across all time windows were then fitted using separate DCMs for each condition. (C) The normalised spectral output of fitted DCMs show near perfect overlap with the observed spectral densities. (D) For each individual, between-condition effects were estimated in a reduced hierarchical model of fluctuations in different subsets of parameters. Across these parametric empirical Bayesian summaries of individual participants, models explaining the spectral changes with fluctuations in time constants have an exceedance probability of > 95%.

**Table 1.** **EEG features of NMDAR-Ab Encephalitis patients**. Patients were selected from routine clinical service based on paroxysms identified on clinical EEG recordings

| *ID* | **Sex** | **Age**<br>**(years)** | **EEG background** | **EEG paroxysms** |
| --- | --- | --- | --- | --- |
| *N001* | M | 2 | Normal | Isolated slow waves |
| *N002* | F | 15 | Normal | Intermittent rhythmic slow, regional left frontal |
| *N003* | F | 9 | Diffusely continuous slow | Intermittent rhythmic slow, generalised, maximum bifrontal |
| *N004* | F | 1 | Normal | Runs of spike and wave complexes, generalised; Isolated sharp waves |
| *N005* | F | 3 | Diffusely continuous slow | Intermittent slow, generalised, maximum bifrontal |
| *N007* | F | 10 | Diffusely continuous slow | Intermittent slow, generalised |
| *N008* | F | 14 | Diffusely continuous slow | Near continuous spike and wave complexes in sleep, generalised, maximum right frontal (ESES) |
| *N009* | F | 11 | Diffuse continuous slow | Intermittent slow, generalised, maximum bifrontal |

Cortical source estimation for the paroxysmal EEG activity was performed and ‘virtual electrode’ responses extracted from the most active sources ^24^. For each patient, DCMs were independently fitted to power spectral density averages of each available condition (e.g. background, short paroxysms, ongoing rhythmic activity, Fig 4B). Individually fitted DCMs (with near perfect model fits) (Fig 4C) were subsequently combined in (within-patient) between-condition hierarchical (PEB) models that explained the condition specific differences with changes in synaptic time constants (**τ**), between-population inhibitory connections (*g_i_*) between-population excitatory connections (*g_e_*), or within population modulatory connections (*g_m_*). Across participants, models explaining spectral differences as arising from differences in time constants offer the best explanation of the virtual electrode data (with an exceedance probability of >95%,Fig 4D).

### NMDAR-Ab alter the response to intrinsic fluctuations in synaptic dynamics

The DCM of human data provides us with an estimate of condition-specific changes in synaptic parameters. We extracted the first principal component of these changes across all conditions and subjects, and applied them to the control and the NMDAR-Ab positive mouse CMC model.

The differences between the parameter estimates from the control and NMDAR-Ab positive model result in different spectral outputs; even when applying the same time constant changes. Overall, the NMDAR-Ab positive context results in higher delta-band power and less high frequency power (Fig 5A-B). Crucially, delta power was higher in the NMDAR-Ab positive model across a wide range of time constant fluctuations (Fig 5C). Furthermore, small changes in the synaptic parameters identified with the patient data are caused large changes in delta power in, and only in, the NMDAR-Ab positive model. This is manifest as low frequency paroxysmal activity, when the synaptic parameters change slightly in the NMDAR-Ab positive model, but not the control (Fig 5D). Technically, this abrupt change in dynamics with a small change in parameters is known as a *phase transition;* suggesting that antibody-positive effects on synaptic coupling move the network towards a critical regime in which small fluctuations in synaptic time constants produce qualitatively different dynamics (i.e., paroxysmal EEG abnormalities).

**Fig 5.**
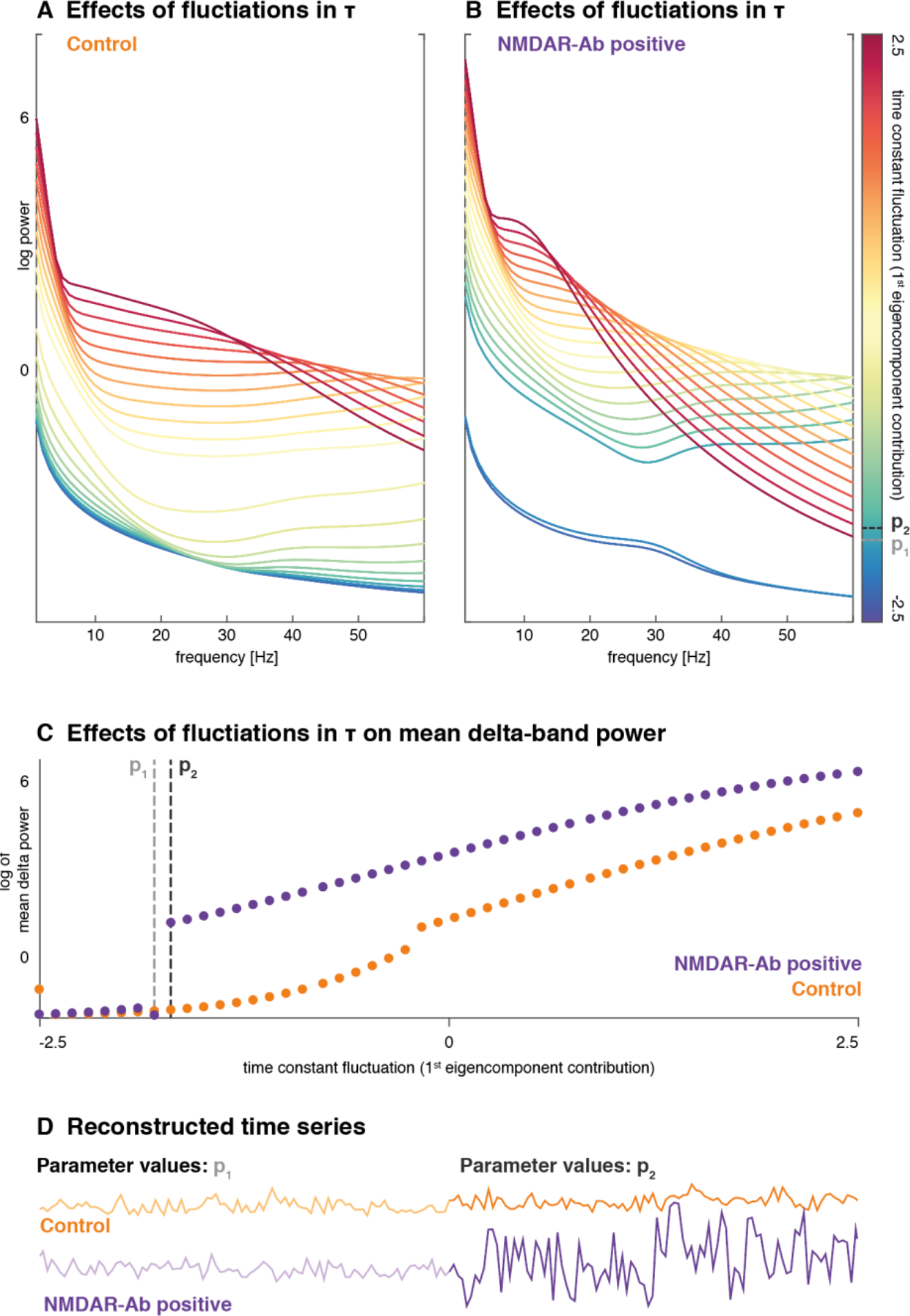
NMDAR-Ab sensitise the microcircuit to intrinsic fluctuations in time constants. Here, we apply a summary component of the time constant fluctuations estimated from human patients to a cortical microcircuit model derived from the control mice (left), and the NMDAR-Ab positive mice (right). (A-B) The same fluctuations cause spectral outputs containing much higher relative delta power in the model estimated from NMDAR-Ab positive mice. (C) This figure shows log of mean delta power for a range of smoothly increasing time constant fluctuations. In the low parameter range (-2.5 to -1.5), there is a large jump in delta power, suggesting that there are two distinct dynamic states separated by small differences in parameter values. (D) Example reconstructions of time series for parameter values at two very close parameter values (*p*_1_and *p*_2_) are shown for control and NMDAR-Ab positive models. The sudden increase in delta power is visible as paroxysmal change in the time series in the NMDAR-Ab positive context, whilst the control time series appear continuous. This sudden change in dynamics with a small change in parameter space is known as a phase transition.

## Discussion

The study presented here reveals common synaptic mechanisms underlying a range of electrophysiological disturbances associated with NMDAR-Ab in a mouse model and paediatric patients: NMDAR-Ab cause a shift in cortical synaptic parameters that is associated with increased low frequency oscillations and predisposes to the low slow wave paroxysms seen in the clinical EEG recordings.

### NMDAR-Ab are associated with high-amplitude low-frequency discharges

NMDAR-Ab cause changes in the spectral composition of resting state LFP of the mouse strain tested. These differences are further revealed on additional exposure to PTZ, with a large PTZ-induced increase in mean delta power in the presence of NMDAR-Ab. The increased delta on the LFP in this mouse model is largely due to intermittent rhythmic slowing; without overt epileptic spikes. Previous analysis of seizure events show that NMDAR-Ab also lower the seizure threshold in these mice ^9^, but seizure events fall largely outside the frequency spectrum analysed here. These spectral changes are in keeping with the kind of abnormalities seen in NMDAR-Ab encephalitis, clinically – with background slowing with or without additional slow wave paroxysms.

An increase in the power of slow frequency components in an EEG or LFP recording is thought to be associated with increased synchronisation of local cortical firing, which itself is regulated by a range of interacting cortical and subcortical systems (e.g. thalamocortical loops ^25, 26^, brain stem monoamine arousal systems ^27^ and intrinsic cortical effects such as astrocytic regulation of synaptic function ^28^). Firing synchrony can occur physiologically (e.g. during sleep), can be associated with non-specific cortical dysfunction (e.g. in the context of an encephalopathy), or be a component of epileptic discharges (apparent in slow-wave component in spike-wave discharges) ^29^.

Synchrony by definition is an emergent feature of population dynamics, rather than a property of any single neuron, but an increase in cortical synchrony may arise from a whole range of different neuronal coupling changes. Many of these can be captured in models that specifically model cortical microcircuitry at the *mesoscale*; i.e. as neuronal ensembles ^21,30,31^. The DCM approach adopted here uses this mesoscale modelling to identify the changes underlying the emergence of hypersynchronous slow wave activity, in the context of NMDAR-Ab.

### NMDAR-Ab cause laminar specific changes in cortical dynamics

Dynamic causal modelling rests on neural mass modelling of coupled neuronal oscillators that are described using specific synaptic parameters (e.g. connection strengths, time constants, activation parameters) and that broadly resemble the laminar structure of the cortex. The neural mass model of a single electromagnetic source contains two pairs of coupled neuronal oscillators that support slower (deep oscillator: deep pyramidal cells, inhibitory interneurons) and faster (superficial oscillator: superficial pyramidal cells, spiny stellate cells) activity ^32^. These population dynamics modelling cortical system (in this case, a single cortical column), with individual parameters that exert highly non-linear effects on the system is output. The parameterisation of these models is rooted in biophysical properties of individual neurons, but describe average characteristics of populations of functionally related neurons; i.e., composite properties emerging from the features of individual cells.

At this mesoscale, PTZ and NMDAR-Ab produce synergistic effects that result in excessive synchrony not seen in other experimental conditions. Our results suggest that increases in low frequency power can be explained by a combination of: (1) an increase in superficial cortical excitatory coupling, largely associated with PTZ exposure, and (2) opposing changes in the dynamics of the superficial oscillator pair (spiny stellate and superficial pyramidal cells, Fig 3).

The changes in synaptic dynamics align time constants in a gradient along the CMC coupling chain, with the slowest time constants in the deep pyramidal cells, and fastest time constants in the superficial pyramidal cells – and gradual steps between. This reduces the stepwise difference in time constants along the CMC chain compared to the standard CMC configuration. This parameterisation allows a dominant frequency to resonate across – and recruit – the whole column, thus producing the high amplitude slow frequency patterns observed. Thus interestingly, slow wave activity appears to be under the control of the faster, superficial oscillator pair in the CMC model, with both NMDAR-Ab and PTZ having profound and relatively specific effects on their dynamics. This is in keeping observations from invasive recordings of slow wave activity in human patients with epilepsy, which implicate superficial cortical coupling in the regulation of slow wave sleep activity ^33^.

### Different molecular changes show converging effects at the neuronal population level

The synaptic parameters of the CMC model employed in DCM are population summaries of a variety of cellular effects, encompassing emergent properties and multiple nonlinearities ^34^. Time constants at the population level are essentially descriptions of the dynamics of post-synaptic integration affected by multiple factors, such as background firing frequency, membrane conductance, intra- and extracellular ion composition, and the dynamics of receptor types present in the membrane to name but a few ^35^. Connection strengths at the population level summarise the effect one population has over another, and may include effects mediated via subpopulations within (e.g. self-connections are assumed to be mediated via local inhibitory interneuronal populations). Because a number of different effects may converge on the same population parameters – and individual molecular effects may only be expressed in certain conditions – the link between molecular change and population parameters is nontrivial.

Exposure to NMDAR-Ab has been reported to cause a number of changes in the postsynaptic glutamate response, including a reduction in overall postsynaptic potential, a reduction in late postsynaptic currents, and a faster return to baseline ^10, 11^. In intact neuronal circuits, NMDAR exert differential control over excitatory and inhibitory populations, leaving the populations differentially affected by NMDAR-blockade ^16, 36^.

PTZ is believed to act as an antagonist to γ-Aminobutyric acid type A (GABA-A) receptors by directly blocking ionophores ^37^. GABA-A receptors are fast inhibitory receptors with a wide-spread, region and cell-type specific set of post-synaptic effects ^38^. These include inhibitory post-synaptic potentials, but also inhibition of dendritic excitatory post synaptic potentials via extrasynaptic GABA-A receptors, which is particularly pronounced at the cortical pyramidal cells ^39^. In some neuronal cell types and at certain developmental stages GABA-A can cause excitatory post synaptic potentials ^40^, and GABA transmission can exert direct or indirect control over excitatory NMDAR-dependent synaptic transmission ^41, 42^.

With this range of different cellular effects, it is unlikely one can capture the breadth of NMDAR-Ab and PTZ related effects in a small subset of population model parameters. However, the effects on delta-band power can be reproduced well with a few principal components; comprising largely just two main effects: (1) decreasing the time constants of superficial pyramidal cells relative to excitatory spiny stellate cells, and (2) increasing the excitatory coupling between spiny stellate and superficial pyramidal cells.

There are a number of possible and convergent changes at the molecular level associated with NMDAR-Ab and PTZ exposure that could explain these population level effects. The time constant changes in superficial pyramidal cells may result from being switched towards (faster) AMPA mediated excitatory inputs (due to the NMDAR-Ab mediated internalisation of NMDAR) and a change in membrane conductivity (due to PTZ-mediated blocking of extrasynaptic GABA-A receptors). The change in excitatory connection, on the other hand, is consistent with a disinhibition of excitatory EPSPs under GABA-A blockade with PTZ (i.e. a block of so-called shunting inhibition) ^39^.

### NMDAR-Ab sensitise the cortical column to spontaneous paroxysmal EEG

In the patients with NMDAR-Ab encephalitis, there is no experimental control over NMDAR-Ab exposure, and our sample of patients is heterogeneous, representative of clinical practice (e.g. age, gender, timing of EEG, timing of initial diagnosis, etc). Moreover, these patients show a diverse range of paroxysmal, short-term changes in EEG dynamic patterns that are visually apparent, allowing us to probe spontaneous fluctuations of DCM parameters that may underlie discrete pathological brain states.

Patient-specific modelling, as facilitated by DCM, allows inference on patient-specific parameters in a generic model of the cortical column. Thus applying DCM analysis to this diverse sample, one can access two types of results: (1) Qualitative: i.e., identify the parameters whose changes underlie the dynamic abnormalities seen in EEG; (2) Quantitative: i.e., establish the numerical range of parameter fluctuations that can be applied to other specified DCMs.

Surprisingly, and despite the variety of EEG abnormalities described, consistently across patients, models with changes in time constants – i.e. synaptic transmission dynamics – best explained the observed transitions between background activity and paroxysms. Furthermore, we could summarise changes in these parameters along a single (principal component) axis. We used this component to enforce similar fluctuations in the fully specified DCMs derived from the mouse model analysis; asking whether the baseline context (i.e. the parameterisation derived from NMDAR-Ab positive or control animals) alters the impact of parameter changes of the magnitude observed in human patients.

Indeed the dynamic responses of the two types of models are very different: In the context of NMDAR-Ab, overall greater delta-band power is observed, and there are regimes of parameter space that contain boundaries between very different dynamic states ^43^. This structural instability underwrites phase transitions of the sort seen in seizure activity. In the control parameterisation, the same changes have a much less pronounced effect, and do not induce overt slow wave paroxysms. In short, it appears that paroxysmal EEG activity in patients may be best explained by *normal* fluctuations in synaptic time constants that occur in an *abnormal* regime of synaptic parameter space.

Overall, these findings provide integrative evidence from human patients and a mouse model of NMDAR-Ab encephalitis suggesting that (1) NMDAR-Ab cause electrophysiological abnormalities via a small number of synaptic changes, which may lend themselves to targeted therapeutic interventions; e.g., by exploiting laminar and/or cell-type specific effects of transcranial current stimulation. ^44^ And (2) paroxysmal abnormalities can be explained by persistent baseline changes that render cortical microcircuitry particularly sensitive to (potentially normal) fluctuations in synaptic coupling. Future research may reveal whether similar approaches have diagnostic value when performed on patient EEGs alone.

## Limitations

The modelling approach presented here allows unique insights into possible mechanisms underlying empirically observed phenomena. Although DCM has been applied to a wide variety of neurophysiological studies – and its validity has been assessed repeatedly ^19,20,45^ – there are certain limitations to the approach adopted here.

First, the modelling can only be applied to existing data – this places restrictions on study design (e.g. pre-NMDAR-Ab exposure EEGs are not usually available from patients) and limit the approach to a subset of testable hypotheses. Second, like all inference, DCM is based on specific assumptions regarding the underlying neuronal architecture – all activity presented here is presumed to emerge from microcircuitry consistent with the CMC model, and only given this assumptions can we estimate the parameters and evidence for or against specific model configurations.

Most importantly, we have reduced a complex brain-wide pathology of interacting systems to changes in a cortical microcircuit. Thus, we are ignoring interactions between different cortical regions, as well as the influence of subcortical structures, such as thalamus and brain stem, which (especially in the context of encephalopathy and slow wave abnormalities) will exert a powerful influence over cortical states. Although these effects can be accommodated in the model as random effects, they are not modelled explicitly.

## Acknowledgments

We are grateful to the patients and their families for agreeing to contribute to this study.

## Methods

### Collection and classical analysis of mouse LFP

The mouse model and associated procedures have been previously described ^9^. Briefly, plasma with NMDAR-Ab (Immunoglobulin G, IgG) was obtained with informed consent from three female NMDAR-Ab positive patients with neuropsychiatric features, movement disorder and reduced level of consciousness. Control IgG was purified from serum from two healthy individuals. C57BL/6 female mice aged 8–10 weeks were housed and examined according to ARRIVE guidelines and all analyses were performed with the observer blinded to injected antibody.

Wireless telemetry transmitters (Subcutaneous transmitter A3028B-CC from Open Source Instruments Inc) were implanted in a subcutaneous pocket over the right flank. Two craniotomies were performed at 1mm lateral and 1mm caudal from bregma. Electrode screws were fixed into the drilled holes with dental cement. After a five-day monitored recovery period, eight microliters of purified IgG (patient, or control) was injected slowly into the left lateral ventricle through a single additional craniotomy made 1mm left lateral and 0.45mm caudal from bregma.

Mice were housed in a Faraday cage during wireless LFP data collection. To test seizure susceptibility, 40mg/kg of PTZ was given intra-peritoneally and the mice were observed for 45 minutes following injection. The 45-minute time period immediately preceding PTZ injection was used as control segment.

Raw LFP data was analysed in Matlab. Sliding-window (30s windows, 15s steps) Fourier estimates of power over frequency were used to statistically compare the different conditions. ANOVA over mean delta-band power (1-4Hz) was used to estimate the effects of the two main interventions (NMDAR-Ab, PTZ) and their interaction on LFP signal composition.

### DCM analysis of mouse LFPs

Modelling of the mouse LFP recordings can be divided into the following steps (summarised in Fig 6). Dynamic causal modelling was performed using SPM12, an academic software package (www.fil.ion.ucl.ac.uk/spm). All analysis code and raw data are available online (https://github.com/roschkoenig/NMDAR_Encephalitis, requires Matlab 2014b or later and SPM12).

**Fig 6.**
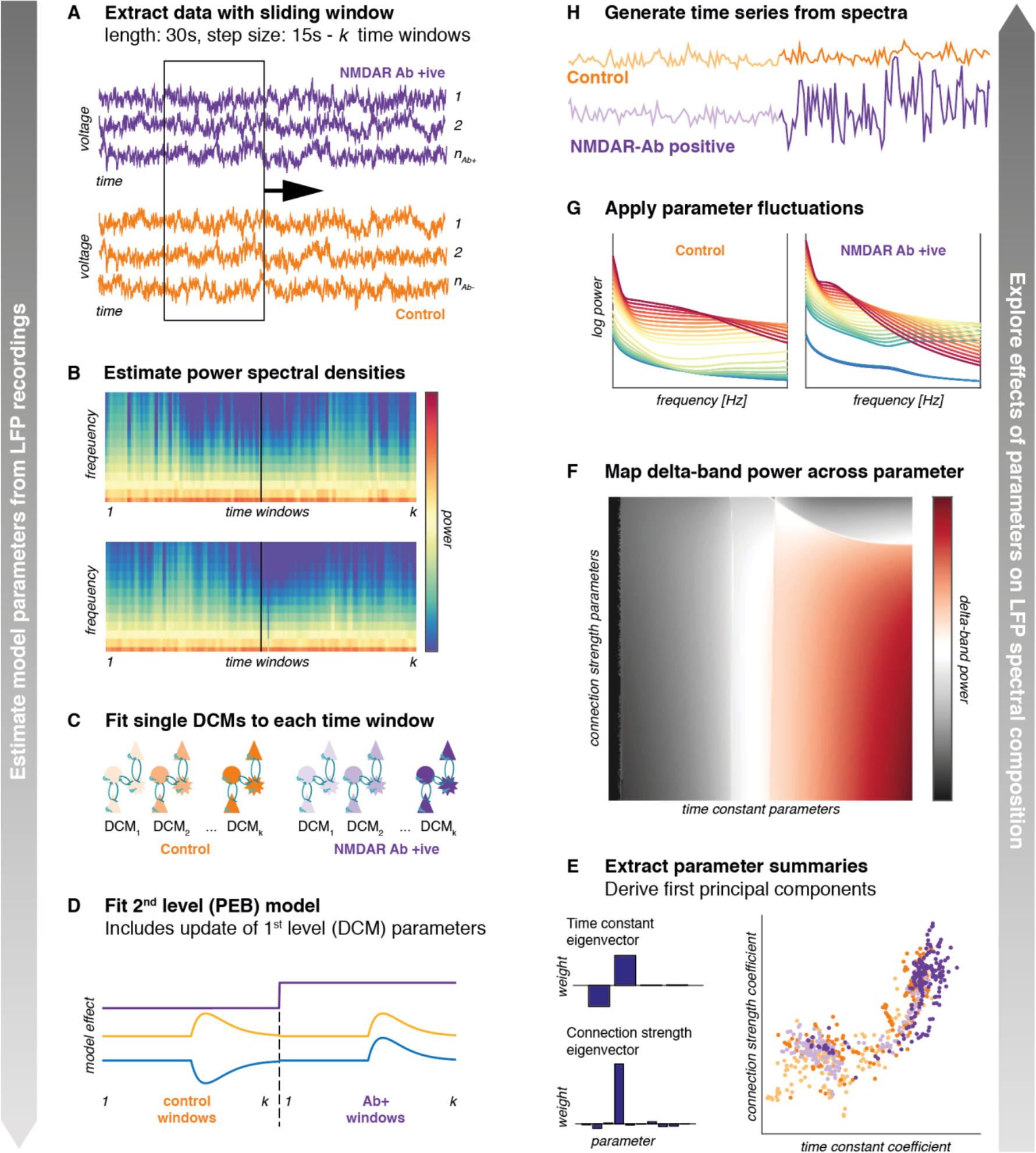
Modelling approach to mouse LFP recordings. Modelling was designed to extract relevant parameters (left hand panels), and then explore the effects of those on delta power (right hand panels). (A) For both pre- and post PTZ injections, 45 minutes of LFP recordings were extracted for each mouse. A sliding window was used to extract a sequence of time windows for further analysis. (B) Power spectral densities were estimated for each time window, which are the basis for the DCM model fit. (C) Single-source DCMs comprising a single CMC model were fitted to each time window separately. (D) Using a parametric empirical Bayesian approach to fit a second level between-DCM general linear model we extracted parameter variations explained by specific experimental effects, and updated first level DCM parameters. (E) From the updated first level DCMs, we extracted all parameters and summarised them in two principal components over time constants and connection strengths, retaining the first component summaries of the fitted DCMs. (D) Starting from the baseline model specification, we applied the reduced (i.e. first principal component) summaries of the parameter changes to simulating cross spectral outputs of the neural populations, yielding a map of delta-power across the ensuing two dimensional parameter space. (G) We then applied quantitative parameter changes observed in patient EEGs (summarised as their first principal component) to the control, and NMDAR-Ab baseline model specifications to explore the effects of parametric fluctuations on spectral output. (H) To further illustrate the effects of parametric fluctuations, we applied and inverse Fourier transform to generate substitute time series – illustrating the nature of the changes in a time trace.

1. Inversion of separate single-source DCM for each time window (performed on group-average data)
2. Second level (PEB) model to explain parameter changes over time, based on experimental interventions
3. Forward modelling to explore effects of parameter changes on specific output measures (e.g. delta power)

Individual time windows were assumed to be relatively stationary within the 30s sliding time window in line with previous DCM analyses of EEG seizure activity ^23, 46^. Each time window was modelled as originating from a single cortical source comprising four coupled neuronal populations (i.e. a single cortical column modelled as single CMC). DCM employs a standard variational Laplace scheme to fit the parameters of a specified neural mass model to empirical data ^18^, whilst also providing a (free energy) measure of the Bayesian model evidence. The combination of posterior parameter estimates and free energy subsequently allows for computationally efficient modelling of group effects across individual DCMs, further exploited with the PEB analysis ^22^.

A second level model (PEB) was used to estimate parameter changes associated with the experimental modulations. Specifically, each time window was associated with a numerical value representing the absence or presence of NMDAR-Ab (0 or 1), the estimated PTZ concentration (range 0 to 1, modelled as first order kinetics after intraperitoneal injection), and an interaction term (range -1 to 1). PEB employs Bayesian model reduction based on the effects specified model parameters, effectively modelling between-window changes in parameter as a mixture of random effects and systematic modulation of each parameter by the main effects provided in the PEB model specification. Thus, inversion at the second (between-window) level provides posterior parameter estimates for first level model parameters (i.e. neuronal physiology) that are associated with second level parameters (i.e. experimental modulation) across the whole series of individual DCMs.

The DCMs are fully specified models of spontaneous neuronal activity and can therefore be used to explore individual parametric effects on overall spectral output. Here, we utilise the parameter estimates derived as the group mean in the PEB analysis as baseline. We then extract the first principal components of time constant, and connection strength variations across all individual time window DCMs (Fig 6E), providing a summary of covarying changes in parameters that explain most of the variance among samples. We then systematically vary the contribution of each of these two components in 300 discrete steps each around the baseline estimates. This yields 300*300 = 90,000 parameterisations for a single source DCM, and for each of these the spectral output can be estimated. We can use this to visualise scalar output measures (e.g. log mean delta band power) across a section of a two-dimensional parameter space (Fig 6F). This combines the benefits of fitting generative (i.e. forward) models to empirical data (i.e. data features to model parameters), and exploring the effects of specific parameters on model output through forward modelling (i.e. model parameters to data features ^47, 48^)

In a last step, we apply the first principal component of the parameter variations in time constants derived from the patient sample NMDAR-Ab positive and control baseline parameterisations of the DCM, to explore the condition specific responses of the cortical column parameterisation to fluctuations in the time constant parameters. We then use and inverse Fourier analysis to illustrate the sort of paroxysmal responses that would be expected based on the spectral predictions under specific parameter combinations(Fig 6H)

### Patient selection and EEG recording

Patients were selected from routine clinical service at a tertiary paediatric specialist hospital that is a regional referral centre for patients with presumed autoimmune encephalitis. Patients were selected based on (1) symptoms consistent with autoimmune encephalitis, (2) positive laboratory testing for NMDAR-Ab at some point during their clinical course, (3) availability of routine clinical EEG recording during the acute phase of their illness, (4) presence of visually apparent EEG abnormalities. Anonymised clinical information was provided by the patients’ care team with written, informed consent provided by the patients’ legal guardians. All patients met the Graus criteria for a clinical diagnosis of NMDAR-Ab encephalitis ^49^.

All EEGs used in this analysis were standard clinical recordings (21 electrodes, International 10-20 Electrode Layout, 30 min recording time, 256 Hz sampling frequency, 1-70 Hz digital Butterworth bandpass filter). EEGs were visually analysed by two clinicians with expertise in EEG interpretation (RER, GC), identifying paroxysmal abnormalities, as well as segments of artefact free awake background EEG that were used for further analysis.

### DCM analysis of patient EEG paroxysms

EEG analysis was designed to identify mechanisms underlying the frequently observed paroxysmal abnormalities in patients with NMDAR-Ab encephalitis. The purpose of this modelling approach is to identify a small set of parameters that can explain the transition between background activity and EEG paroxysms for each individual patient. The analysis can broadly be summarised as follows (also shown in Fig 7).

**Fig 7.**
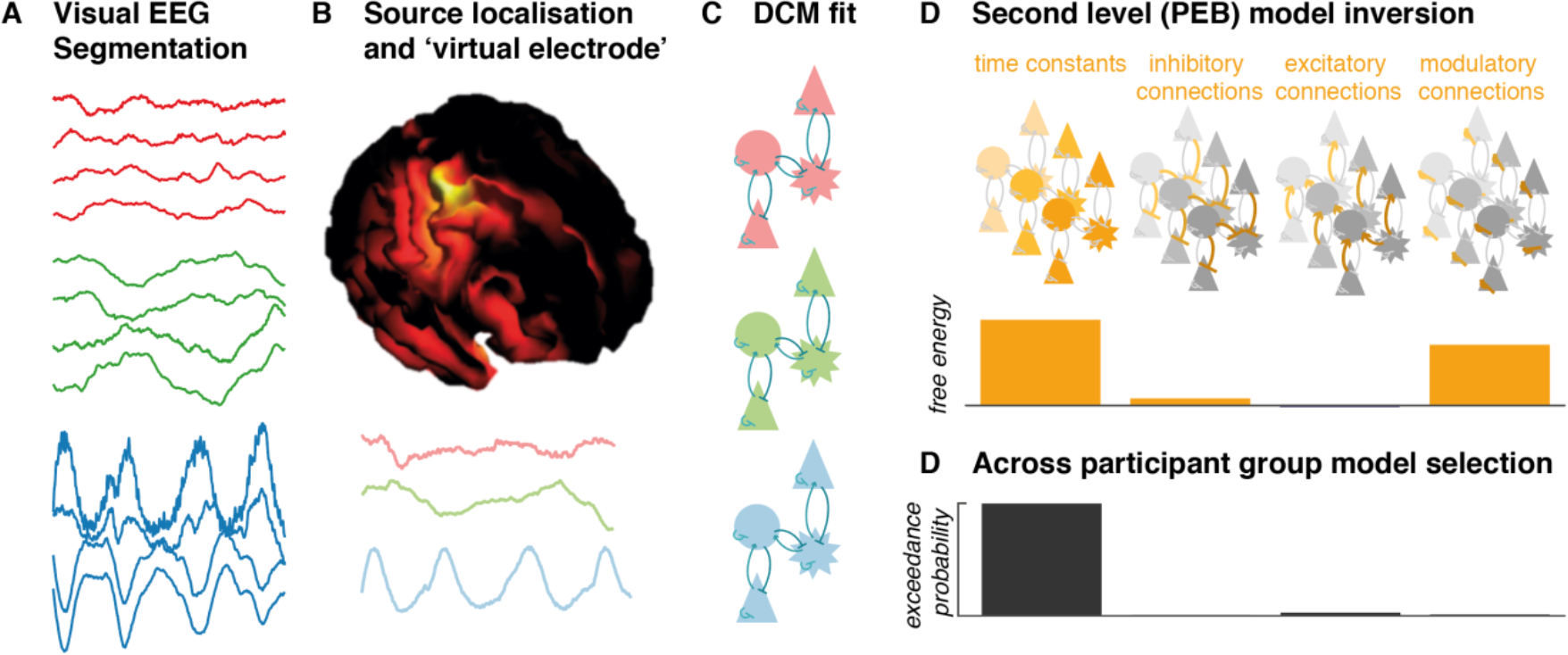
DCM analysis approach for patient EEG recordings. (A) Visual analysis was performed to identify segments of artefact free background EEG, as well as visually apparent paroxysms of abnormal activity (which were further separated into isolated and rhythmic abnormal activity). (B) This activity was source localised using an IID approach. Subsequent modelling was performed using a virtual electrode estimate of LFP activity at the identified source (bottom of this panel). (C) Single source DCMs comprising a single CMC were fitted separately to power spectral density averages of background, and paroxysmal activities (D) PEB was employed to reduce within subject differences between individual DCMs to specific subsets of parameters. The model space was designed to distinguish between sets of models where time constant, inhibitory connections, excitatory connections, or modulatory connections explained variations among conditions. (D) A random effects Bayesian model comparison between these alternative PEB models helped identify which parameters best explain the fluctuations across the whole group of subjects.

1. Visual identification of paroxysmal and background EEG activity source localisation and ‘virtual electrode’ source wave form extraction
2. Fitting of single source DCM to each ‘virtual electrode’ summary of paroxysmal and background data
3. Inversion of hierarchical (PEB) model explaining all within-subject EEG patterns Bayesian model comparison between sets of reduced models at the group level (random effects analysis)

Patients were selected on clinical EEG with reported dynamic abnormalities (ranging from evidence of mild encephalopathy to overt epileptiform activity). EEGs were reviewed by two clinicians with EEG experience (RER, GC) and segments containing normal awake background, as well as paroxysmal abnormalities (isolated slow waves, intermittent rhythmic slow activity, and overt epileptiform activity) identified. Paroxysmal activity was averaged across visually identified 2s windows and source localised using an IID (independent and identically distributed) approach in SPM12 ^50^. At the cortical location with maximal activity, a single virtual electrode trace was extracted for each of the paroxysmal and background activity windows and used for further DCM analysis ^46^.

This ‘virtual LFP’ activity was modelled using a single CMC source. An average of all paroxysm time windows, and all background time windows was inverted separately, producing 2–3 fully specified DCMs per subject. These were subsequently combined into a single hierarchical (PEB) model for each patient, in which only a subset of specific parameters were allowed to vary. A model space was created at the level of these second level models were either time constants, inhibitory between-population connections, excitatory between-population connections, or inhibitory self modulatory connections were allowed to vary to explain the difference between paroxysms and background activity (see Tab 2). Random effects Bayesian model comparison across these second level models uses the approximation to model evidence from the variational Laplace model inversion (i.e. the free energy) to compare the evidence for any given model parameterisation, given the empirical data ^19^.

**Table 2:**
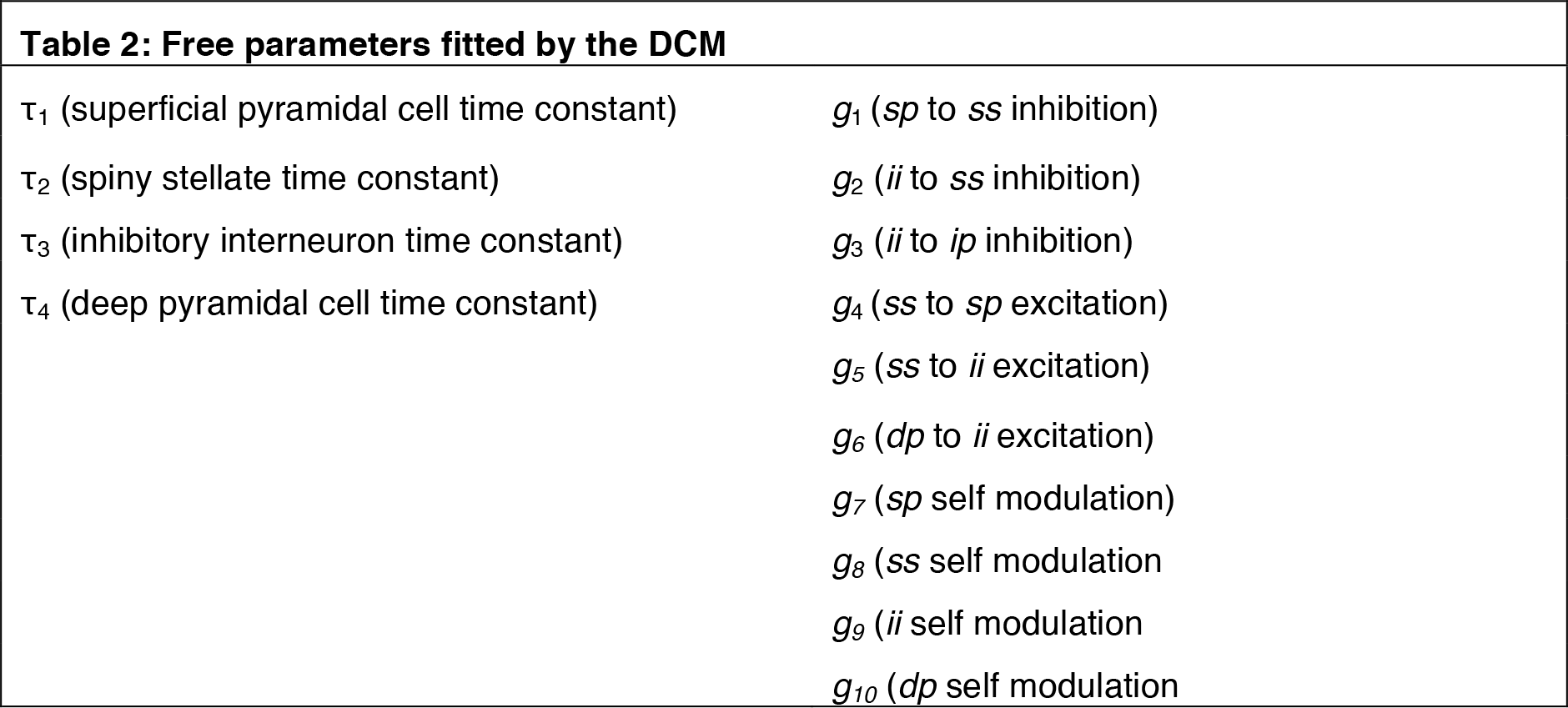
Free parameters fitted by the DCM

